# Microbial Gladiators: Unraveling the dynamics of carbon substrate competition among heterotrophic microbes

**DOI:** 10.1101/2023.09.19.558456

**Authors:** Samuel M. McNichol, Fernando Sanchez-Quete, Stephanie K. Loeb, Andreas Teske, Sunita R. Shah Walter, Nagissa Mahmoudi

## Abstract

Growing evidence suggests that interactions among heterotrophic microbes influence the efficiency and rate of organic matter turnover. These interactions are dynamic and shaped by the composition and availability of resources in their surrounding environment. Heterotrophic microbes inhabiting marine environments often encounter fluctuations in the quality and quantity of carbon inputs, ranging from simple sugars to large, complex compounds. Here, we experimentally tested how the chemical complexity of carbon substrates affects competition and growth dynamics between two heterotrophic marine isolates. We tracked cell density using species-specific PCR assays and measured rates of microbial CO_2_ production along with associated isotopic signatures (^13^C and ^14^C) to quantify the impact of these interactions on organic matter remineralization. The observed cell densities revealed substrate-driven interactions: one species exhibited a competitive advantage and quickly outgrew the other when incubated with a labile compound while both species seemed to coexist harmoniously in the presence of more complex organic matter. Rates of CO_2_ respiration revealed that co-incubation of these isolates enhanced organic matter turnover, sometimes by nearly twofold, compared to their incubation as mono-cultures. Isotopic signatures of respired CO_2_ indicated that co-incubation resulted in a greater remineralization of macromolecular organic matter. These results demonstrate that simple substrates promote competition while high substrate complexity reduces competitiveness and promotes the partitioning of degradative activities into distinct niches, facilitating coordinated utilization of the carbon pool. Taken together, this study yields new insight into how the quality of organic matter plays a pivotal role in determining microbial interactions within marine environments.

## Introduction

Marine environments harbor diverse heterotrophic microbes that are critical for the transformation and remineralization of organic matter [1–3]. Much of this organic matter consists primarily of complex compounds, necessitating the use of extracellular enzymes to hydrolyze them into subunits that are small enough to be imported into the cell [4, 5]. However, not all heterotrophic microbes possess the necessary enzymatic machinery for degrading complex organic matter. Some microbes rely on consuming simpler compounds that can be directly taken up and metabolized. This leads to a trophic cascade, wherein primary degraders with the ability to secrete hydrolytic enzymes initiate the breakdown of complex compounds, while secondary degraders rely on metabolic by-products and/or degradation products as growth substrates [6–9]. Throughout this process, a complex web of interactions emerges that encompass both competition and cooperation [10, 11]. These interactions can influence the composition and diversity of microbial communities and can have cascading effects on ecosystem functioning and biogeochemical cycling [12, 13].

Microbial interactions are inherently dynamic, continuously shaped by their surrounding environment and the intricate interplay among microbes [14, 15]. The competitive exclusion principle of ecology, often referred to as Gause’s Law, asserts that in stable environmental conditions, when two species compete for the same resource, even a slight advantage will eventually result in the dominance of the better competitor, leading to the extinction of the inferior species or its adaptation to a new, non-competitive niche [16]. This principle has been extensively validated through experiments involving simple substrates (i.e., small molecules that can be directly transported into the cell) [17, 18]. However, our understanding of how complex mixtures of carbon substrates influence species dynamics remains limited, leaving uncertainties about their impact on the turnover of environmental organic carbon on both short and long timescales. Interestingly, recent studies suggest that the chemical complexity of carbon substrates affects the way bacteria interact [19, 20]. Complex substrates appear to reduce competition for resources and increase cooperative interactions among primary degraders while labile substrates promote competition. It is not entirely clear why, but one possible explanation is that complex substrates promote niche differentiation that minimizes direct competition and ultimately leads to coordinated degradative activities [21].

Microbes inhabiting marine environments often encounter fluctuations in both the quality and quantity of carbon inputs, ranging from simple sugars to complex, high-molecular weight compounds [22, 23]. These fluctuations have the potential to alter interactions among heterotrophic microbes and shift dynamics from functional redundancy (competition) to functional complementarity (cooperation) [9, 24]. Consequently, it is crucial to investigate these interactions within the context of natural mixtures of carbon substrates, as opposed to a single substrate. However, the inherent complexity of natural organic matter makes it challenging to differentiate between carbon pools that support the growth of heterotrophic microbes and those that do not using traditional microbiological or geochemical approaches. Natural abundance carbon isotopic measurements (^13^C and ^14^C) have emerged as a powerful analytical tool to overcome these challenges and gain valuable insights into the origin and age of the organic matter utilized by microbes for growth and respiration [25, 26]. When heterotrophic organisms break down organic matter, the respired CO_2_ retains the same Δ^14^C signatures and nearly identical δ^13^C signatures as the carbon sources. Therefore, the ^13^C and ^14^C content of respired CO_2_ can serve as a proxy for the source organic matter [27].

We previously carried out comparative incubations using a novel bioreactor system (Isotopic Carbon Respirometer-Bioreactor; IsoCaRB, [28]) to explore organic matter remineralization in marine sediments by two different heterotrophic marine isolates (i.e., primary degraders) [29]. This system allows us to continuously measure respiratory CO_2_ production and its associated isotopic (Δ^14^C, δ^13^C) signatures, providing us with a comprehensive picture of the rate, quantity, and type of organic matter remineralized by each species. Recognizing that interactions among microbes can influence the transformation of natural organic matter, our goal in this study was to examine how competition affects the growth dynamics of these primary degraders and to quantify the impact of these interactions on organic matter turnover. Here, we conducted competition experiments using the IsoCaRB system with these same two primary degraders (*Vibrio splendidus* 1A01 and *Pseudoalteromonas sp.* 3D05). These experiments were carried out with carbon substrates of varying chemical complexity, thereby allowing us to directly test the impact of substrate complexity on interactions between these marine heterotrophs. Additionally, this unique experimental approach allowed us to contextualize these interactions within a biogeochemical framework, which enabled us to establish a quantitative link between competition (or its absence) and the rate and extent of organic matter remineralization.

## Materials and Methods

### Bacterial strains and culture conditions

For our competition experiments, we used two marine strains (*Vibrio sp.* 1A01 and *Pseudoalteromonas sp.* 3D05) that were previously isolated from coastal ocean water samples (Canoe Beach, Nahant, MA) for their ability to degrade complex substrates [7, 12] (Table S2). Both strains were grown concurrently under identical growth conditions to ensure that they would reach similar cell densities at the same time. The strains were inoculated from frozen glycerol stocks into 25 ml of liquid Marine Broth 2216 (Difco #279110; Franklin Lakes, NJ, USA) in a 125 ml Erlenmeyer flask and incubated overnight at room temperature with shaking at 145 rpm until reaching log-phase. Subsequently, cells were transferred (1% inoculation) into 50 ml of Tibbles-Rawlings (T-R) minimal medium (see Table S3 for detailed recipe, [30]) that was supplemented with 0.5% (w/v) D-(+)-glucosamine hydrochloride (TCI America, #G0044; Portland, OR, USA) as a carbon source. Cell density was monitored by measuring optical density (OD) at 600 nm, based on a calibration curve between OD and colony forming units (CFUs), to ensure that the starting concentration of cells for the co-culture experiments was equivalent to that of the mono-culture experiments conducted by Mahmoudi et al. (2020) [29]. Once cells reached a desired cell density of 5 × 10^8^/ml, 50 ml of cell culture was harvested by centrifugation for 10 min at 3000 xg (Beckman Coulter Allegra X30-R Centrifuge; Montreal, QC, Canada) and washed two times with T-R minimal medium containing no carbon source. The cell pellets from each isolate were then resuspended in 1 ml of T-R minimal medium containing no carbon source and injected into the IsoCaRB system simultaneously using a 3 ml syringe (BD Biosciences #309657; Franklin Lakes, NJ, USA) and a 20-gauge needle (Sigma-Aldrich #Z192511; St. Louis, MO, USA).

### Co-culture experiments in the IsoCaRB system

Co-culture experiments were conducted in the IsoCaRB system which allows for continuous monitoring and collection of microbially respired CO_2_ for δ^13^C and Δ^14^C isotopic analyses. These time-resolved isotopic measurements provide information on the source and age of natural organic matter being utilized, as well as its rate of remineralization. The IsoCaRB system is comprised of a gas delivery and purification system, a custom Pyrex culture vessel, an inline CO_2_ detector and integrated LabVIEW data-logging program, custom CO_2_ traps, and a vacuum extraction line. The system is first purged of residual atmospheric CO_2_ by sparging with CO_2_-free helium for at least 24 h. Gas flow is then changed to 100 ml min^-1^ (20% O_2_ in helium) approximately 1 h prior to injection of bacterial cells. Respiratory CO_2_ is carried to an online infrared CO_2_ analyzer (Sable Systems CA-10; Las Vegas, NV, USA) where concentrations are quantified in real time and continuously logged to a desktop PC using a custom LabVIEW program (National Instruments; Austin, TX, USA). This CO_2_ is continuously collected as successive fractions in custom molecular sieve traps. Subsequently, CO_2_ is recovered from the traps by baking (530°C for 30 min) under vacuum within 24 h of collection, then cryogenically purified, quantified, and stored in flame-sealed Pyrex tubes. Each experiment is allowed to proceed until CO_2_ concentrations resume near-baseline values. Gaseous CO_2_ concentration measurements are converted to a rate of CO_2_ generation per unit volume of growth medium (µg C L^-1^ min^-1^). Lastly, the culture vessel is equipped with a sampling port which permits direct sampling of the medium throughout the incubation. Details regarding the standard operating procedure for the IsoCaRB system, including sterilization and assembly, sample preparation, and CO_2_ collection and purification are described in Beaupré et al. (2016) [28].

All experiments in the IsoCaRB system were carried out under aerobic conditions (∼20% O_2_) to ensure an abundant supply of oxygen. In contrast, organic carbon was limited to a total of 880 mg in each experiment to foster competition among primary degraders. To test the influence of substrate type on competition dynamics, co-culture experiments were carried out using three distinct sources of organic carbon, each varying in complexity (Fig. S1). A simple sugar (glucosamine) was used to evaluate competition for readily metabolizable substrate that could be directly taken up into cells without the requirement of extracellular enzymes. Natural marine sediments collected from Guaymas Basin were utilized for experiments to examine competition for more complex carbon substrates found in natural mixtures. Guaymas Basin is a young, active spreading center in the central Gulf of California where sediments are impacted by hydrothermal heating [31]. This leads to the production of more labile compounds, such as petroleum hydrocarbons [32], and smaller organic acids [33–35]. Conversely, cool sediments in off-axis regions do not undergo heating and organic matter found in these areas resembles that of typical marine sediment. Therefore, co-culture experiments were conducted using both hydrothermally-influenced sediments, which encompassed a mixture of labile and complex substrates, as well as unimpacted, cool sediments comprised primarily of complex substrates. For the experiments conducted with sediments, microbially respired CO_2_ was trapped and collected for ^13^C and ^14^C analysis to assess how competition impacted the source and age of the organic matter remineralized.

#### A) Glucosamine Hydrochloride

*Vibrio sp.* 1A01 and *Pseudoalteromonas sp.* 3D05 were co-incubated with 2.2 g (equivalent to ∼880 mg of C) of filter-sterilized D-(+)-glucosamine hydrochloride (TCI America, #G0044) along with 2 l of T-R minimal medium in replicate. The slurry was subsampled in replicate every 12 h to track the number of cells; two 10 ml subsamples of slurry were collected and stored at -80°C for subsequent DNA extraction and cell quantification.

#### B) Sediments

A total of three competition experiments were performed with sediments: two with unimpacted sediment and one with hydrothermally-influenced sediment. Experiments were carried out with the same sediment samples previously used by Mahmoudi et al. (2020) (Table S4) [29]. In addition, the exact same mass of sediment was used in the competition incubations as the mono-culture incubation to allow us to assess organic matter remineralization of each isolate in the absence of competition. Sediment cores were retrieved during dives with the research submersible HOV *Alvin* (Woods Hole Oceanographic Institution) on a cruise to the Guaymas Basin in December 2016. Cool, unimpacted sediments had an in-situ temperature of 3-5°C as measured by the *Alvin* Heatflow probe, matching the bottom water temperature. Hydrothermal sediments ranged from 3°C at the seawater interface to 60-90°C at 20 cm depth. Additional details regarding sediment core characteristics, collection, and sediment preparation are described in Mahmoudi et al. (2020) [29]. Briefly, the upper 0-10 cm of sediment cores were homogenized and titrated to pH 2-2.5 with 10% HCl in an ice bath to remove carbonate species. Subsequently, homogenized sediments were freeze dried and sterilized by gamma-irradiation via a ^137^Cs (radioactive cesium) source to receive a total dose ∼40 kGy prior to use in experiments.

*Vibrio sp.* 1A01 and *Pseudoalteromonas sp.* 3D05 were co-incubated at room temperature with approximately 22 g (dry weight) of decarbonated, sterilized sediment. Sediment total organic carbon (TOC) content was previously measured to be 4% [29], equating to ∼880 mg of organic carbon (OC) used for each incubation experiment. The sediment slurry was subsampled for cell quantification (via dPCR) every 5-12 h, with sampling occurring more frequently at the beginning of the experiment. Microbially respired CO_2_ was continuously collected as successive fractions in custom molecular sieve traps. The duration of each CO_2_ fraction ranged from 5-28 h to ensure that no fraction contained more than 2 mg of carbon. Microbially respired CO_2_ was recovered from the traps by baking (530°C for 30 min) under vacuum and subsequently cryogenically purified, quantified, and flame-sealed in Pyrex tubes for isotopic analysis.

### Isotopic analysis of microbially respired CO_2_

CO_2_ fractions that were collected during the co-culture experiments with sediment were sent to the National Ocean Sciences Accelerator Mass Spectrometry (NOSAMS) Facility at the Woods Hole Oceanographic Institution for Δ^14^C and δ^13^C analysis. An aliquot of each sample was split for ^13^C measurement, with the remainder reduced to graphite [36] for ^14^C measurement by accelerator mass spectrometry (AMS). All isotopic data are corrected for background contamination associated with the IsoCaRB system as described in Mahmoudi et al. (2020) [29]. Stable isotope values (^13^C) are reported versus the VPDB standard. Radiocarbon values (^14^C) are reported in Δ^14^C notation, where Δ^14^C is the relative deviation from the ^14^C/^12^C ratio of the atmosphere in 1950 corrected for kinetic fractionation using measured δ^13^C values [37].

### Apportioning the contribution of carbon pools to microbial respiration

To evaluate the contribution of different carbon pools to microbial respiration, we employed a mass balance model, recognizing the diverse array of sedimentary organic matter available for microbial utilization [38, 39]. Mass balance models are valuable tools for determining the sources of substrates for biogeochemical processes and are often employed to decipher the mixing of pools within a sample based on measured isotopic values. Given that we used the exact same sediment samples as Mahmoudi et al. (2020) [29], we applied a similar mass balance model and presumed three major carbon sources: (1) phytoplankton-derived compounds, (2) microbially-produced acetate and/or other fermentation products, and (3) pre-aged organic carbon (OC). For hydrothermal sediments, we presumed that the pre-aged OC endmember is predominantly composed of hydrothermally derived petroleum compounds resulting from magmatic heating processes [40]. Conversely, in unimpacted sediments, this older carbon pool is assumed to be primarily composed of aged terrigenous organic matter derived from land plants [41, 42]. Taking these assumptions into account, we used this mass balance model to predict the fractional utilization of all three sources for the competition experiments with hydrothermal and unimpacted sediment, including which pools were preferentially respired at different times of the incubation:

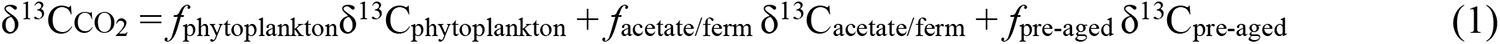

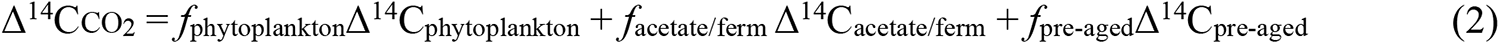

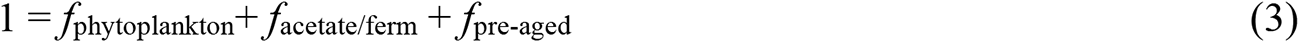

where δ^13^CCO_2_ and Δ^14^CCO_2_ correspond to the measured isotopic values of the respired CO_2_. Fractional contributions (*f’*s) and their uncertainties were estimated as the means and standard deviations of solutions to Equations 1 – 3, which were solved by Monte Carlo resampling (10,000 times) with normally-distributed pseudo-random noise added to each isotope measurement (Δ^14^C ± 50 ‰; δ^13^C ± 2 ‰) and end-member isotopic signature. More information regarding the assumptions and associated uncertainties for each end-member value can be found in the Supporting Information.

### Quantification of individual bacterial strains

The genomes of *Vibrio* sp. 1A01 and *Pseudoalteromonas* sp. 3D05 were previously sequenced, assembled and deposited in project PRJNA478695 (Table S2) [12]. Ten candidate short-amplicon (∼180 bp) primer sets targeting nonoverlapping regions of the genome for each strain was developed using Geneious bioinformatics and primer design software (version 11.1.5, Biomatters Ltd.). To validate primer set performance, aliquots of purified DNA were extracted from isolates and amplified, and the PCR product was visualized on a 1.5% agarose gel in 1X TAE for 60 min at 100 V. Assay performance was assessed by running a standard series from 1 × 10^1^ to 1 × 10^6^ genome copies/rxn using SYBR chemistry and analyzing both experimental and *in silico* melt curves (with uMelt, [43]). Based on the reaction efficiency, sensitivity, and *R*^2^, one primer set was selected for each bacterial isolate and subsequently used to track growth dynamics by analyzing subsamples from the competition experiments (Table S5).

DNA was extracted in replicate from subsamples collected during the co-culture experiments using the Sox DNA Isolation Kit (MetagenomBio Inc.; Waterloo, ON, Canada) according to the manufacturer’s protocol. Genome quantification for *Vibrio sp.* 1A01 and *Pseudoalteromonas sp*. 3D05 were performed in a duplex dPCR reaction using the QIAcuity Probe PCR kit and the QIAcuity Digital PCR system (QIAGEN; Hilden, Germany). Concentrations and thermocycling conditions were performed according to the manufacturer’s instructions. Briefly, dPCR reactions of 40 µl composed of 1X Probe PCR Master Mix, 0.8 µM of each primer, 0.4 µM of each probe, and 0.5 ng of DNA template for both *Vibrio sp.* 1A01 and *Pseudoalteromonas sp.* 3D05 were run in a single well using a 24-well nanoplate of 26k partitions and the following conditions: 1 cycle of initial activation at 95°C for 2 minutes, and 40 cycles of denaturation at 95°C for 15 seconds, followed by a combined annealing/extension at 60°C for 30 seconds. Data was analyzed using the QIAcuity Software Suite 2.1.8.20 and the baseline for positive partitions was adjusted based on the negative and positive controls. Final genome copies were reported according to the dilution factor of each subsample.

## Results and Discussion

### Bacterial growth dynamics are influenced by substrate complexity

Growth dynamics, measured as genome copies/µl, varied between the bacterial isolates depending on the substrate that was present (Fig. 1; Fig. S1). During incubation with glucosamine, *Vibrio* sp. 1A01 exhibited rapid growth within the first ∼12 hours, surpassing that of *Pseudoalteromonas* sp. 3D05, indicating a dominance of *Vibrio* sp. 1A01 cells. Conversely, when both isolates were incubated with both hydrothermal and unimpacted sediment, *Pseudoalteromonas* sp. 3D05 cell numbers exceeded those of *Vibrio* sp. 1A01 (Fig. 1B, 1C). However, this growth pattern exhibited nuances influenced by the complexity of the carbon substrates. During incubation with hydrothermal sediment, which contains low-molecular weight (MW) compounds that do not require external processing using extracellular enzymes [35], *Pseudoalteromonas* sp. 3D05 cell numbers were substantially higher than *Vibrio* sp. 1A01. In the case of unimpacted sediment, *Pseudoalteromonas* sp. 3D05 cell densities were only slightly higher than *Vibrio* sp. 1A01 throughout the experiment, with minimal changes over time. These patterns in cell densities were found to be reproducible during replicate incubations with both glucosamine and Guaymas Basin sediment (Fig. S2).

**Figure 1.**
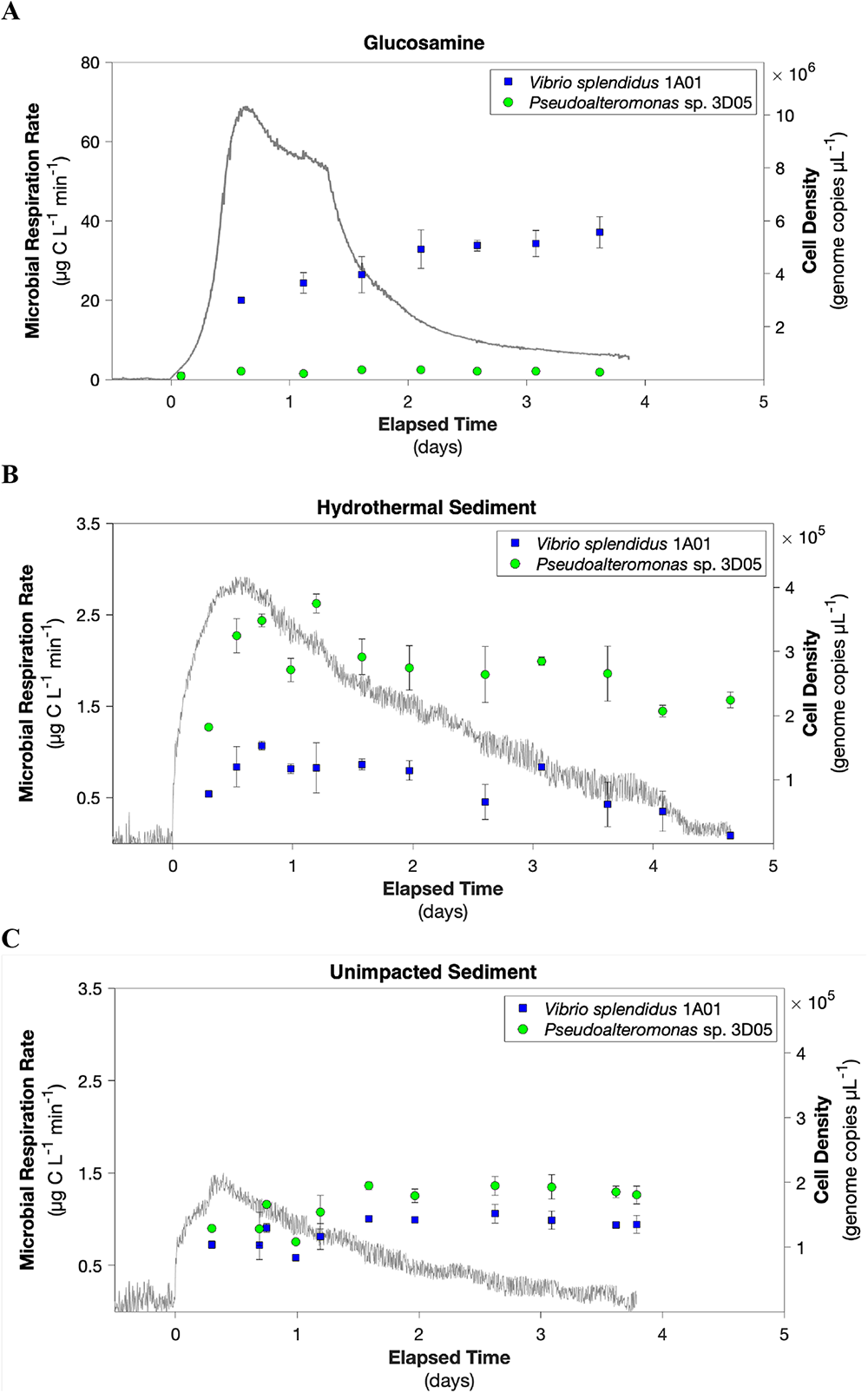
Microbial CO_2_ respiration rates (gray line) and cell densities (squares and circles) measured during co-culture incubations of *Vibrio splendidus* 1A01 and *Pseudoalteromonas* sp. 3D05 with (A) glucosamine; (B) hydrothermal Guaymas Basin sediment; and (C) unimpacted Guaymas Basin sediment. Error bars represent the standard error between dPCR measurements from replicate subsamples.

The chemical complexity of carbon substrates has the potential to exert diverse effects upon the ecology and interactions among microbes [44–46]. Our data reveal that substrate complexity influences the growth dynamics and interactions between primary degraders. When both species are grown with a simple substrate, *Vibrio sp.* 1A01 outcompetes *Pseudoalteromonas* sp. 3D05 and becomes the dominant species throughout the incubation period. In contrast, *Pseudoalteromonas* sp. 3D05 appears to outgrow *Vibrio sp.* 1A01 in the presence of more complex substrates that require extracellular enzymes. Our observed trends may stem in part from differences in the metabolic potential between these two marine heterotrophs. Previous genomic analysis revealed *Pseudoalteromonas sp.* 3D05 possesses approximately four times more gene copies for extracellular peptidases (Table S2), providing a competitive edge for degrading complex substrates. However, the dominance of *Pseudoalteromonas* sp. 3D05 is greatly reduced in the presence of the most complex mixture of carbon substrates, specifically in the presence of unimpacted sediments. It is hypothesized that complex substrates have the potential to foster cooperative interactions and drive the separation of degradative activities into specialized niches [21, 47]. Conversely, when a labile substrate that can be easily transported into the cell is present, competition may intensify as cooperation might not confer any advantage. This can result in the selection of microbial species that are more adept at acquiring and utilizing resources efficiently.

Our findings corroborate prior studies that have suggested complex carbon substrates can increase the potential for niche complementarity thereby reducing competition among microbes. Experimental work using nine different lignocellulolytic bacterial isolates observed that when three-species mixed cultures were grown in media enriched with lignocellulose there was synergistic growth and positive interactions. However, when the same mixed cultures were grown in the presence of glucose as the sole carbon source, there appeared to be competition and negative interactions, resulting in lower cell densities [19]. Likewise, microcosm experiments that constructed aquatic microbial communities of varying levels of species richness from a pool of 16 bacterial isolates found that microbial interactions were strongly attenuated by changes in carbon substrates over time [48]. During the early succession when labile substrates were abundant, strong negative interactions were observed within bacterial communities. As succession progressed and more recalcitrant substrates were utilized, a shift toward more neutral interactions was observed.

### Co-incubation of primary degraders leads to enhanced organic matter turnover

We observed a consistent respiration pattern when bacterial isolates were co-incubated with both simple and complex substrates. Upon the introduction of cells, there was a rapid increase in CO_2_ production, peaking within the first 12 to 24 h, then decreasing to near-constant values within 4-5 days across all experiments (Fig. 1). This ‘single-peak’ respiration pattern aligned with the observations made when these isolates were incubated individually as mono-cultures [29]. Similar to the observed cell densities, the rate of microbial CO_2_ production between replicate incubations was highly reproducible during incubation with both simple and complex substrates (Fig. S2). The highest rates of CO_2_ production were observed during co-culture experiments with glucosamine which had a maximum CO_2_ production rate of 69 µg C L^-1^ min^-1^ (Fig. 1A). This is not surprising since this substrate can be readily taken up by both isolates without requiring extracellular hydrolysis. In contrast, rates of CO_2_ production were considerably lower during incubation with sedimentary organic matter. However, the CO_2_ production rate was substantially higher during incubation with hydrothermal sediment compared to unimpacted sediment, reaching a maximum rate of 2.9 µg C L^-1^ min^-1^ and 1.4 µg C L^-1^ min^-1^, respectively (Fig. 1B, 1C). Higher respiration rates were also observed when isolates were incubated as mono-cultures with hydrothermal sediment compared to unimpacted sediment [29]. These findings are consistent with the fact that hydrothermal sediment contains more labile, low-MW carbon substrates that can be readily consumed and remineralized to CO_2_ [35].

To investigate the impact of interactions on organic matter turnover, we compared the total amount of respired carbon observed during our co-culture incubations to the amount observed when the isolates were incubated as mono-cultures with these same sediments ([29]; Fig. 2). Co-cultures resulted in higher respiration rates and remineralization of organic matter, as evidenced by the greater quantity of respired carbon that was measured. When both isolates were incubated with hydrothermal sediment, the total amount of respired carbon reached 15.1 mg, more than twice the combined sum of respired carbon during the mono-culture incubations of *Vibrio* sp. 1A01 (2.3 mg) and *Pseudoalteromonas* sp. 3D05 (4.6 mg). Similarly, the total carbon respired during the competition experiments with unimpacted sediment (5.9 mg) was slightly higher than the total sum of carbon respired by *Vibrio* sp. 1A01 (1.9 mg) and *Pseudoalteromonas* sp. 3D05 (3.3 mg) during the mono-culture incubations.

**Figure 2.**
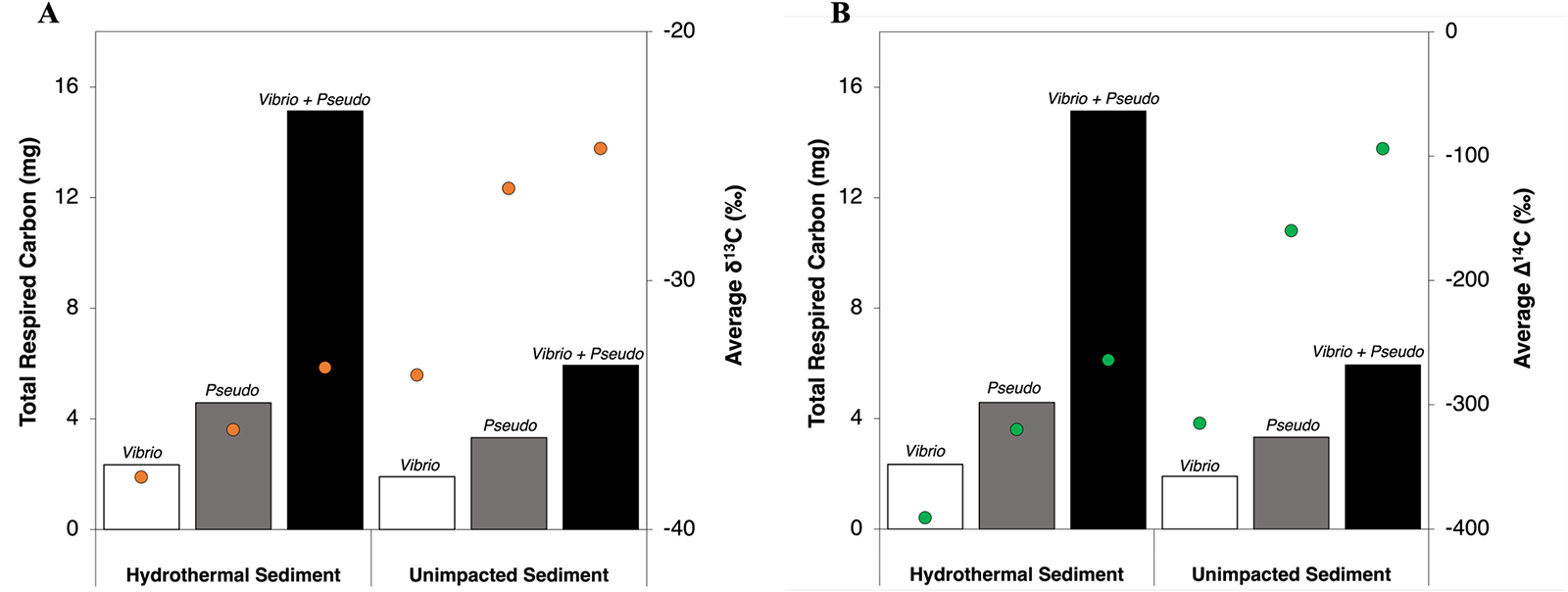
Total mass of carbon respired as CO_2_ (bars) and mass-weighted average δ^13^C (orange circles) and Δ^14^C (green circles) during mono-culture and co-culture incubations of *Vibrio splendidus* 1A01 and *Pseudoalteromonas* sp. 3D05 with hydrothermal and unimpacted Guaymas Basin sediment. Total respired CO_2_ was determined by manometric quantification of microbially respired CO_2_ collected during mono-and co-culture incubation experiments. In order to obtain a mass-weighted average δ^13^C and Δ^14^C value, the isotopic value for each CO_2_ fraction was multiplied by the relative mass of that fraction. These values were then summed over all the fractions within a single incubation.

Our results suggest that when both isolates are grown together in the presence of a complex mixture of substrates, such as sedimentary organic matter, they appear to exhibit varied degradative activities, effectively differentiating into distinct niches. For instance, *Vibrio* sp. 1A01 may rapidly consume labile substrates, compelling *Pseudoalteromonas* sp. 3D05 to utilize more complex substrates that require the secretion of degradative enzymes. This simultaneous utilization of different carbon pools leads to increased organic matter remineralization. Likewise, prior studies have shown that microbial consortia consisting of multiple species show improved capabilities compared to monocultures for the degradation of complex feedstocks such as lignin and lignocellulose [49, 50]. While the exact mechanism that underlies this phenomenon is unclear, it may arise from the specialized nature of degradative enzymes, which target specific bonds, distinct structural regions and/or molecules [5, 51]. As a result, each species may be relatively more efficient at consuming specific carbon pools leading to coordinated degradative activities.

We observed substantially greater organic matter remineralization during co-culture incubation with hydrothermal sediment which contains a larger pool of labile substrates compared to the unimpacted sediment. This pool of readily available substrates may have provided one or both isolates with additional energy to produce extracellular enzymes needed to degrade more complex substrates. This is often proposed as a mechanism for ‘priming’, which is the increased degradation of complex or recalcitrant organic matter resulting from an increase in overall microbial activity due to energy and nutrients gained from the degradation of labile compounds [52]. Although our study did not directly investigate the priming effect, our results suggest that availability of labile compounds could significantly enhance the degradation of more complex organic matter.

### Co-incubation of primary degraders leads to greater remineralization of macromolecular organic matter

To determine the source and age of the organic matter being respired during competition experiments with sedimentary organic matter, a series of CO_2_ fractions were collected successively for natural abundance isotopic analysis (δ^13^C and Δ^14^C). A total of 12 CO_2_ fractions were collected during the co-culture incubation with hydrothermal sediment, while a total of 8 CO_2_ fractions were collected during incubation with unimpacted sediment. Isotopic signatures for one of the CO_2_ fractions (i.e. fraction 4) collected during incubation with the unimpacted sediment was not measured due to sample loss at the radiocarbon facility. However, since this fraction accounted for only ∼5 hours out of a total collection time of 91 hours, we have confidence in our ability to interpret the isotopic trends throughout the incubation in the absence of the isotopic values for this fraction.

The δ^13^C signatures measured during the co-culture experiments showed a progressive enrichment in ^13^C over time, ranging from -37‰ to -24‰ for the hydrothermal sediment, and – 31‰ to -17‰ for the unimpacted sediment (Fig. S3; Table S6). The Δ^14^C signatures observed during co-culture incubation with hydrothermal sediment became consistently more negative over time, ranging from -218‰ to -321‰. However, during the co-culture incubation with unimpacted sediment, the Δ^14^C values started at -111‰ and became more positive in the first ∼24 hours, before becoming more negative (ranging from -94 to -123‰) during the remainder of the incubation (Fig. S3; Table S6).

To understand the dynamic nature of microbial carbon utilization, we employed a mass balance model that enabled us to predict the fractional utilization of available carbon sources including which carbon pools were preferentially respired at different times of the co-culture incubation (Fig. S4, S5). The results of this mass balance revealed that acetate was a primary microbial carbon source during the initial stages of the incubation with both unimpacted and hydrothermal sediment. This observation aligns with expectations that carbon substrates which are relatively labile (e.g., acetate and low-MW organic acids) will be preferentially consumed over more complex substrates [26, 53]. As the incubation progressed, the utilization of acetate diminished over time, giving way to an increased utilization of pre-aged OC from day 1.0 onwards. While pre-aged OC remained a minor source in the case of unimpacted sediment, it became a predominant source of carbon in the incubation with hydrothermal sediment. Interestingly, phytoplankton-derived material was a substantial carbon source throughout the co-culture incubation with both hydrothermal and unimpacted sediment[54].

Our unique experimental approach enabled us to directly test how microbial interactions influenced the remineralization of specific carbon pools during incubation with sedimentary organic matter. We calculated mass-weighted averages of measured δ^13^C and Δ^14^C values for mono-and co-culture experiments to assess how isotopic signatures of respired CO_2_ shifted when both isolates were grown together (Fig. 2). For each incubation, we multiplied the isotopic value for each CO_2_ fraction by the relative mass of that fraction within an incubation (Table S6) and subsequently summed these values to obtain a mass-weighted average δ^13^C and Δ^14^C value (Fig. 2). Mass-weighted average Δ^14^C signatures of respired CO_2_ revealed that co-culture incubations had more positive Δ^14^C values compared to mono-culture incubations, suggesting greater utilization of modern, recently photosynthesized carbon. Similarly, the mass-weighted average δ^13^C signatures of respired CO_2_ during the co-culture incubations was more positive compared to mono-culture incubations, consistent with greater utilization of ^13^C enriched material such as phytoplankton-derived carbon [55]. Next, we assessed the relative contribution of each carbon pool to the total amount of respired CO_2_ in mono- and co-culture experiments (Fig. 3.) This was done by multiplying the estimated end member contributions of the mass balance calculated for each CO_2_ fraction (Fig. S4, S5) by the mass of each fraction (Table S6) and summing the carbon masses. The calculated relative contributions revealed that acetate remained the dominant source of carbon for respiration during both mono- and co-culture incubations. However, there was a substantial increase in the utilization of phytoplankton-derived carbon when both isolates were incubated together compared to when they were incubated individually as mono-cultures. In fact, phytoplankton-derived carbon was estimated to account for 30% ± 3 and 70% ± 5 of all respired CO_2_ during co-culture incubation with hydrothermal and unimpacted sediment, respectively.

**Figure 3.**
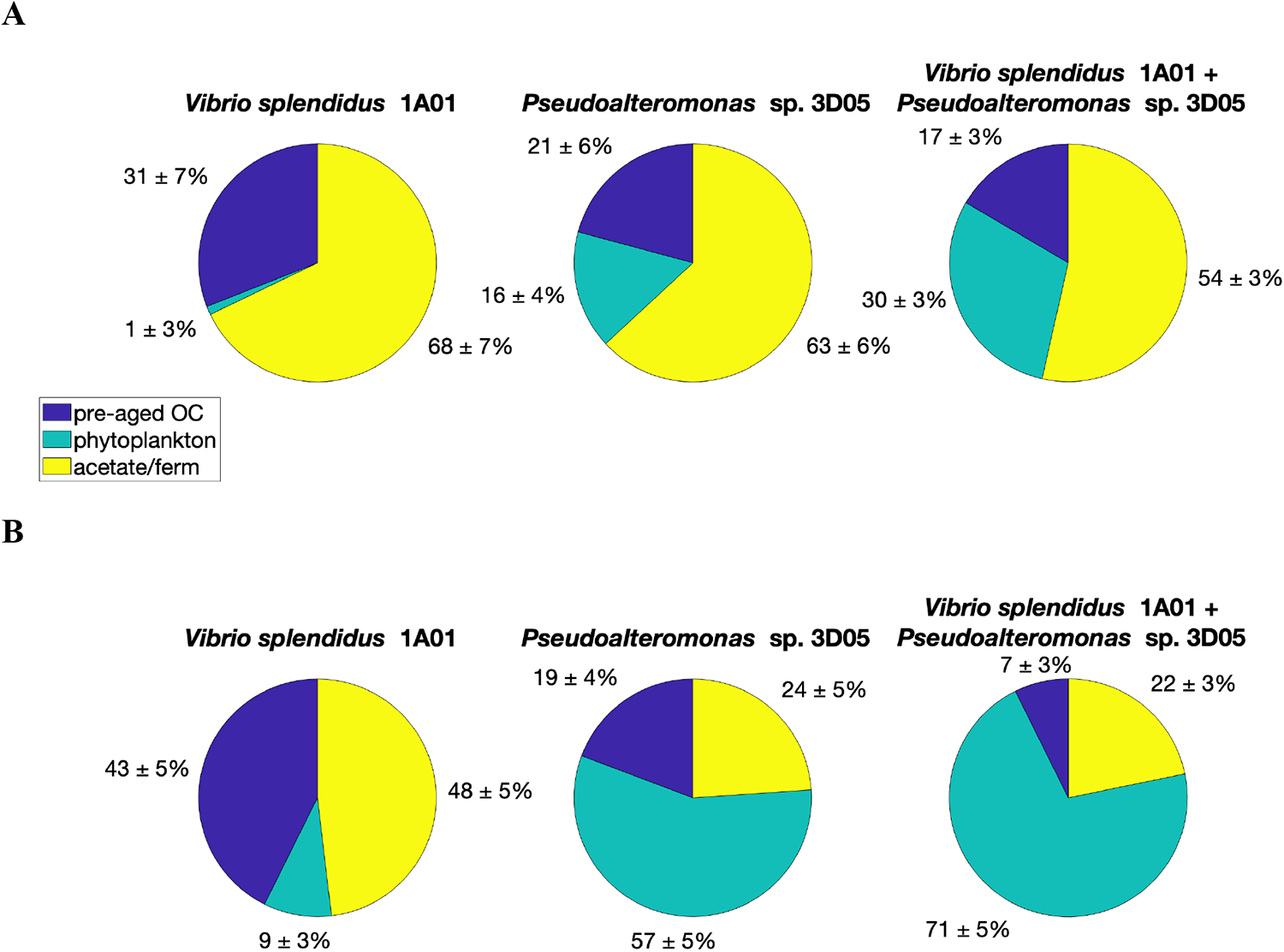
Estimated relative contribution of potential carbon sources to total respired CO_2_ during mono-culture and co-culture incubations of *Vibrio splendidus* 1A01 and *Pseudoalteromonas* sp. 3D05 incubations with (A) hydrothermal and (B) unimpacted Guaymas Basin sediment. Relative contributions were calculated as the relative mass of each fraction attributable to each end member, based on fraction masses and mass balance model output. Percentages and uncertainties of each fraction were estimated as means and standard deviations of solutions to 3 simultaneous mass-balance equations that were solved 10,000 times, in which normally distributed pseudo-random noise was added to each isotope ratio measurement and isotopic signature.

The macromolecular composition of phytoplankton cells (based on dry weight) is dominated by proteins [56]. As a result, phytoplankton-derived carbon pools are assumed to be rich in proteins which require extracellular hydrolysis for consumption. Both *Vibrio sp. 1A01* and *Pseudoalteromonas sp. 3D05* possess multiple gene copies for extracellular peptidases which allows each species to access and utilize this protein-rich carbon pool more readily. The isotopic results reveal that co-culture incubations resulted in greater degradation of carbon pools comprised of larger substrates (i.e. macromolecules), indicative of elevated extracellular enzyme production and activity in the co-culture incubations. It would be expected that the pool of readily available, labile substrates is more rapidly consumed during the co-culture incubations which would prompt one or both species to secrete degradative enzymes earlier in the incubation process. This dynamic interplay would also increase overall turnover of organic matter and the concentration of degradation products and hence the potential for growth of both species. Additional work to assess gene expression may confirm whether the observed increase in organic matter remineralization during the co-culture incubations may be due in part to greater expression of extracellular enzymes by one or potentially both species.

## Conclusion

Uncovering the microbial mechanisms that drive biogeochemical processes in marine environments is essential for unraveling the complexities of the global carbon cycle. There is an increasing number of studies that suggest microbial interactions play a significant role in influencing the efficiency and rate of organic matter turnover [12, 13]. While competition between the two species may arise when both species are grown in the presence of a labile substrate, our results suggest a simultaneous and synergistic utilization of the carbon pool in the presence of complex substrates. These results lend support to the notion that heterogenous complex substrates facilitate a distinct division of labor [21], in which organic matter turnover is enhanced due to coordination of species’ degradative activities. By shedding light on the dynamics of carbon utilization and the consumption of specific carbon pools, this study demonstrates the crucial role of microbial interactions in shaping carbon cycling within marine environments.

## Acknowledgements

This work was supported by grants from U.S. National Science Foundation award (no. 2023656) and Fonds de recherche du Quebec – Nature et technologies (FRQNT) grant (no. 283357). We thank Thi Hao Bui for analytical assistance as well as NOSAMS for isotopic measurements. Targeted on- and off-axis sampling in Guaymas Basin was made possible by NSF grant (no. 1357238) to A.T., and the excellent performance of the *Alvin* and *Sentry* teams during cruise AT37-06).

## Data Availability Statement

Genomes for the isolates used in this study can be found on NCBI under BioProject ID PRJNA478695 under accession numbers PDUR00000000 and PDUS00000000.

