## Supplemental Information for "Microbial Gladiators: Unraveling the dynamics of carbon substrate competition among heterotrophic microbes"

***S1. Mass Balance Modeling***

We used a three-end member mass balance model to estimate the contributions of each carbon pool to microbial respiration during co-culture incubations with hydrothermal and unimpacted Guaymas Basin sediments. This model assumes that there are three major carbon sources within Guaymas Basin sediments: (1) phytoplankton-derived compounds, (2) microbially- produced acetate and/or other fermentation products, and (3) pre-aged organic carbon (OC). Since the co-culture incubations were carried out with aliquots of the exact same homogenized sediment, we presumed that endmembers had similar isotopic signatures as those detailed in Mahmoudi et al., 2020 [1]. Here, we applied the mass balance model and carefully re-assessed the assumed isotopic signatures of endmembers within the context of the isotopic results from the co-culture incubations. In doing so, we slightly adjusted the isotopic values of two end members: δ^13^C value of phytoplankton and Δ^14^C value of acetate and/or fermentation products (see Table S1). We recalculated the estimated contribution of carbon pools to microbial respiration during the mono-culture incubations using these adjusted isotopic values (Figure S4); however, since the adjusted values were within error of the previously set values, we obtained nearly identical estimates as those previously calculated by Mahmoudi et al., 2020 [1]. Below we provide additional details regarding each end member and the assumed isotopic signatures used in this model.

**Table S1.** Isotopic signatures of endmembers used in source apportionment calculations.

|  |  | **δ^13^C (**‰) | **Δ^14^C (**‰) |
| --- | --- | --- | --- |
| Hydrothermal, on-axis, sediments | Acetate | -44 ± 2‰ | -360^2^ ± 50‰ |
|  | phytoplankton | -20^1^ ± 2‰ | -12 ± 50‰ |
|  | pre-aged OC | -23 ± 2‰ | -431 ± 50‰ |
| Unimpacted, off-axis sediments | Acetate | -44 ± 2‰ | -226 ± 50‰ |
|  | phytoplankton | -20^1^ ± 2‰ | -12 ± 50‰ |
|  | pre-aged OC | -24 ± 2‰ | -490 ± 50‰ |

^1^Previously set to -22‰ by Mahmoudi et al., 2020

^2^Previously set to -400‰ by Mahmoudi et al., 2020

*Phytoplankton endmember*

Given that Guaymas Basin contains very productive overlying waters and has a fast sedimentation rate, phytoplankton-derived organic compounds should be a dominant carbon source to seafloor sediments. This material will have an identical carbon isotopic signature in both hydrothermal and unimpacted sediments. The commonly accepted δ^13^C values for phytoplankton-derived carbon ranges from -19 to -23‰. In Mahmoudi 2020 [1], the δ^13^C endmember of phytoplankton was set to -22 ± 1‰ [2]. However, the δ^13^C values measured in our study revealed that this end member was likely closer to the positive end of this range. Thus, we assumed that the δ^13^C value of phytoplankton-derived organic carbon is -20‰ ± 2‰.

The Δ^14^C value of phytoplankton-derived organic carbon is be expected to be consistent with the surface water dissolved inorganic carbon (DIC), since phytoplankton convert DIC to organic matter through photosynthesis. The sediment core encompassed the upper ~10 cm of surface sediment and the sedimentation rate at Guaymas Basin is ~ 1 mm/year [3, 4]; thus, the upper 10 cm of sediment core would include phytoplankton-derived material from 1916 to 2016. The Δ^14^C value of phytoplankton-derived organic carbon was assumed to be -12‰, based on the average Δ^14^C of surface water DIC over this 100-year period.

*Pre-aged OC endmember*

Magmatic heating of thick, organic-rich Guaymas Basin sediments near the axis leads to the active production of petroleum within basin sediments [5, 6]. Although this petroleum is generated at an average sediment depth of ~10 to 50 m, there is vertical migration of these compounds to the sediment-water interface such that surface sediments are rich in petroleum. Therefore, the pre-aged OC pool in hydrothermal sediments was presumed to be dominated by hydrothermal petroleum-derived compounds. The carbon isotopic signatures of Guaymas oils have been previously measured. The δ^13^C endmember of our pre-aged endmember was set to the previously measured value of -23 ± 2‰ [7], while the Δ^14^C endmember was set to the previously measured average value of -431‰ ± 50‰ [8, 9].

Cool, off-axis sediments were collected from unimpacted sampling sites that experienced little to no heatflow (e.g. 5°C below 70cm) and displayed no particular evidence of petroleum generation. Other sources of older carbon to marine sediments can include inputs of terrigenous organic carbon through rivers and streams as well as deep dissolved organic carbon (DOC). The δ^13^C endmember for pre-aged organic carbon was set at -24 ± 2‰, which is consistent with measured δ^13^C values of DOC in the Colorado River which feeds directly into the Gulf of California that ranged from ~ -20 to ‑27‰ [10]. For the Δ^14^C signature, we used the previously assumed value of (-490 ± 50 ‰) that was constrained by Mahmoudi et al., 2020 [1]. This was done repeatedly solving Eq. 1 – 3 with hypothetical pre-aged Δ^14^C signatures (from -500 to -200 ‰ in 5 ‰ increments) using the Δ^14^C values measured during the mono-culture incubations. The ranges of physically-meaningful pre-aged OC Δ^14^C signatures was found to be -500 to -480 ‰ for *Vibrio sp. 1A01* and -500 to -220 ‰ for *Pseudoalteromonas sp. 3D05*. Therefore, the Δ^14^C value of this endmember to lie at the center of this range and prescribed a large uncertainty (-490 ± 50 ‰).

*Acetate and/or other fermentation products endmember*

Microbial acetogenesis is thought to be widespread in marine sediments [11-13]. We assumed that acetate and other low molecular weight acids in Guaymas Basin sediments were primarily produced through heterotrophic acetogenesis which would have resulted in acetate as being ~19.5‰ more depleted relative to δ^13^C_TOC_ (Table S2). Both hydrothermal and unimpacted Guaymas Basin sediments had a δ^13^C_TOC_ value of -24.1‰ and -24.8‰, respectively, thus, in our mass balance model the δ^13^C value of acetate was set to -44 ± 1‰.

The Δ^14^C value of acetate would be assumed to be similar to the initial carbon substrate used by the microbe to produce acetate. However, this initial carbon substrate could potentially be derived from a number of different pools. For the cool, off-axis Guaymas Basin sediments, we could not further constrain the Δ^14^C value of acetate because we had another unconstrained pool (pre-aged OC). Therefore, the Δ^14^C value of acetate in the unimpacted sediment was assumed to have the same value as the bulk organic carbon pool (Δ^14^C_TOC_ = -226‰; Table S2) and the Δ^14^C value of the acetate end member was set to -226 ± 50‰. For the hydrothermal Guaymas Basin sediment, Mahmoudi et al., 2020 [1] had previously constrained the acetate Δ^14^C signatures to 400 ± 50‰. This was done by repeatedly solving Eq. 1 – 3 with hypothetical acetate Δ^14^C signatures (from -450 to +100 ‰ in 5 ‰ increments). However, when we applied this value to the Δ^14^C signatures measured during co-culture incubations with hydrothermal sediment it did not lead to physically-meaningful values. Since both mono- and co-culture incubations were carried out with aliquots of the exact same homogenized sediment, it would be improbable for the end member of this carbon pool to be different. Thus, we re-assessed the acetate Δ^14^C signature by repeatedly solving Eq. 1 – 3 with hypothetical acetate Δ^14^C signatures (from -450 to +100 ‰ in 5 ‰ increments) using the Δ^14^C values measured during both the mono- and co-culture incubations. We constrained the Δ^14^C value of this endmember to a slightly more positive range and led us to set the acetate Δ^14^C signature to 360 ± 50‰.

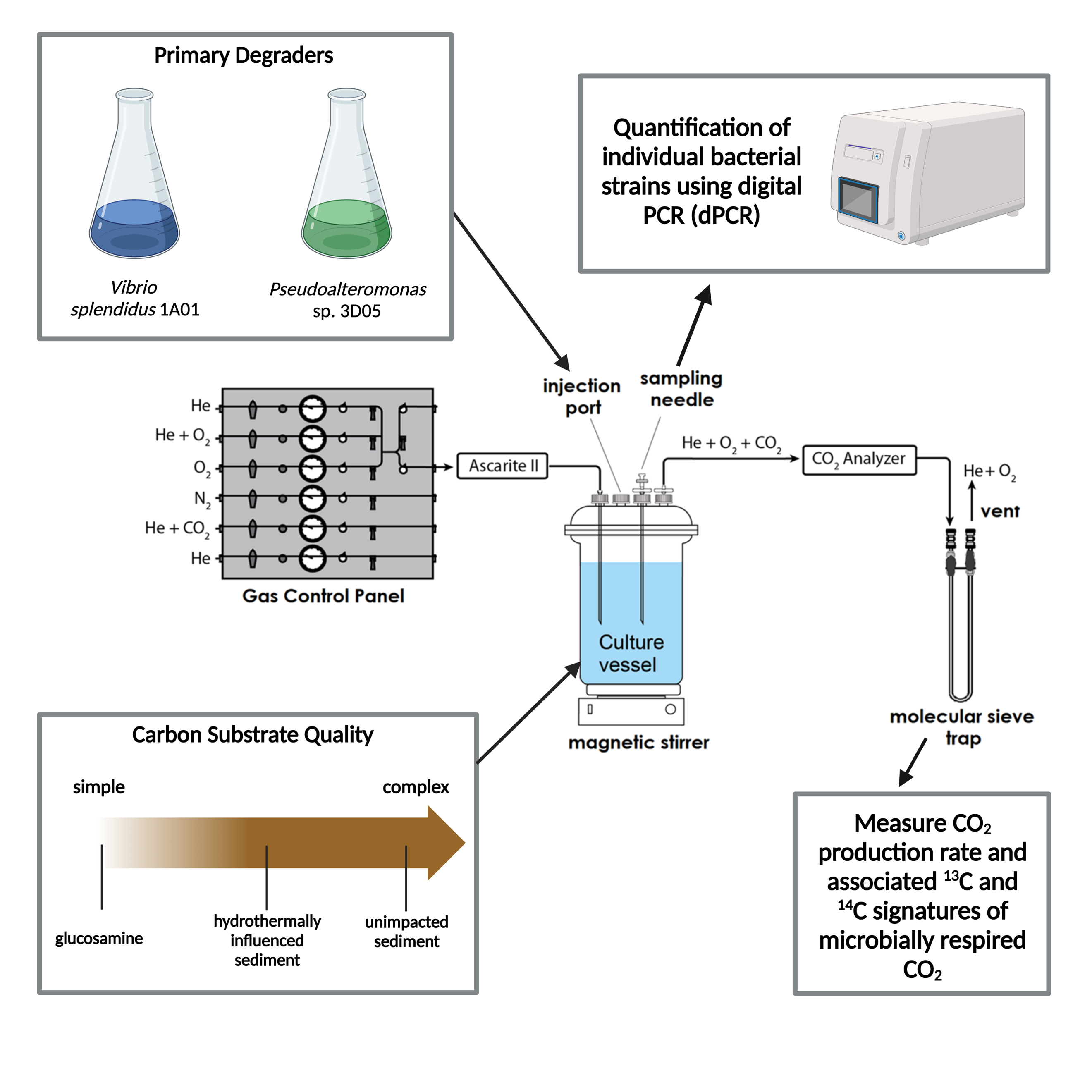

**Figure S1.** Simplified diagram for co-culture competition experiments using the IsoCaRB bioreactor system. *Vibrio splendidus* 1A01 and *Pseudoalteromonas sp.* 3D05 were grown separately to identical cell densities before simultaneous injection into the IsoCaRB culture vessel. The chemical composition of the carbon substrate added to the IsoCaRB system varied from simple to complex. Subsamples were collected from the culture vessel for cell quantification using dPCR, while the respired CO­_2_ was simultaneously collected in successive fractions for δ^13^C and Δ^14^C analysis.

**A**

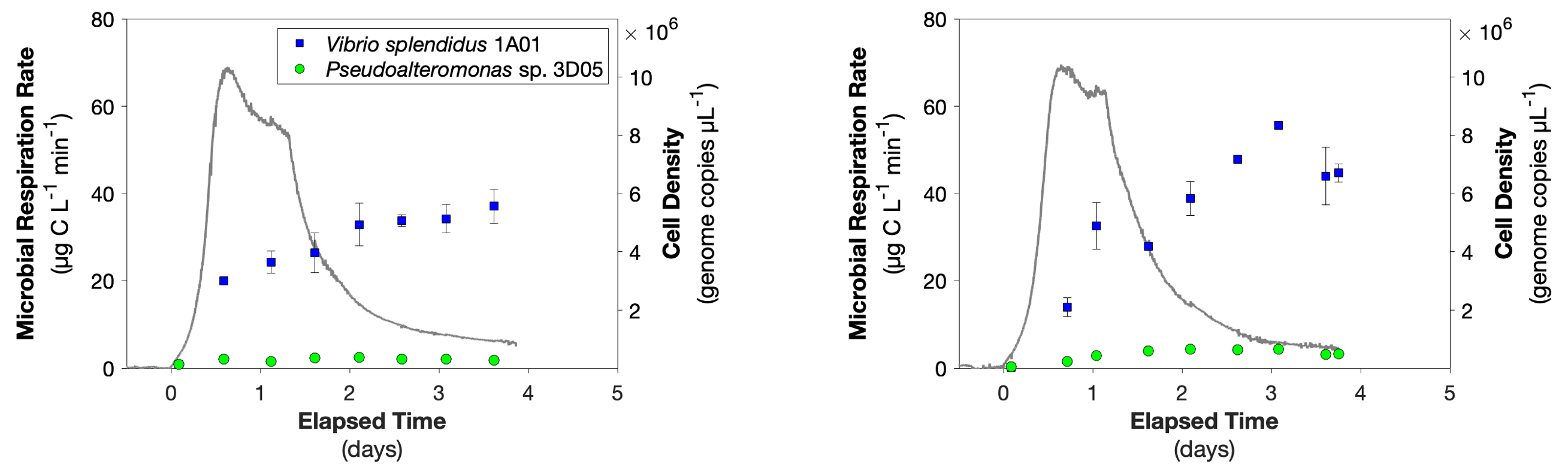

**B**

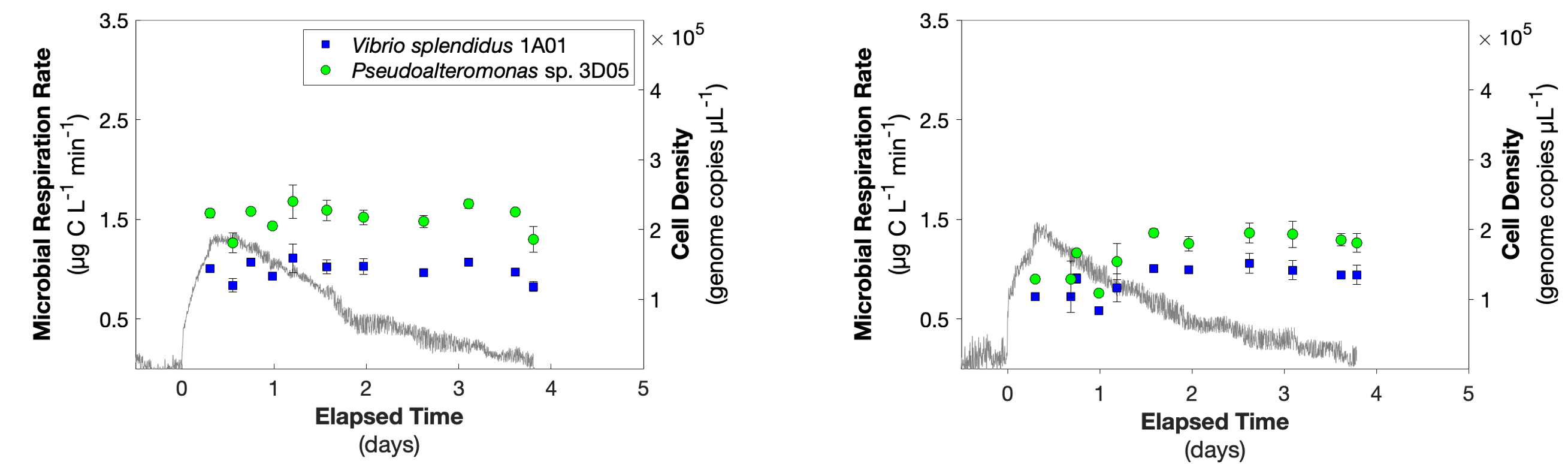

**Figure S2.** Microbial CO_2_ production rates (grey line) and cell density of *Vibrio splendidus* 1A01 and *Pseudoalteromonas* sp. 3D05 during replicate co-culture incubations with (A) glucosamine and (B) unimpacted Guaymas Basin sediment.

**A**

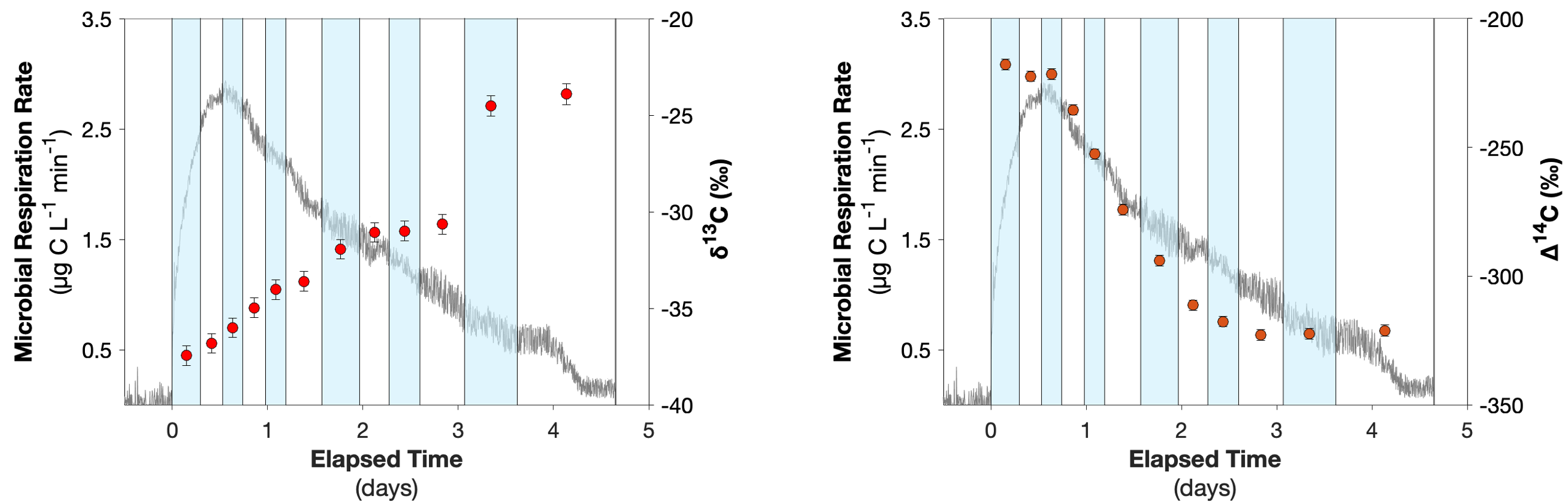

**B**

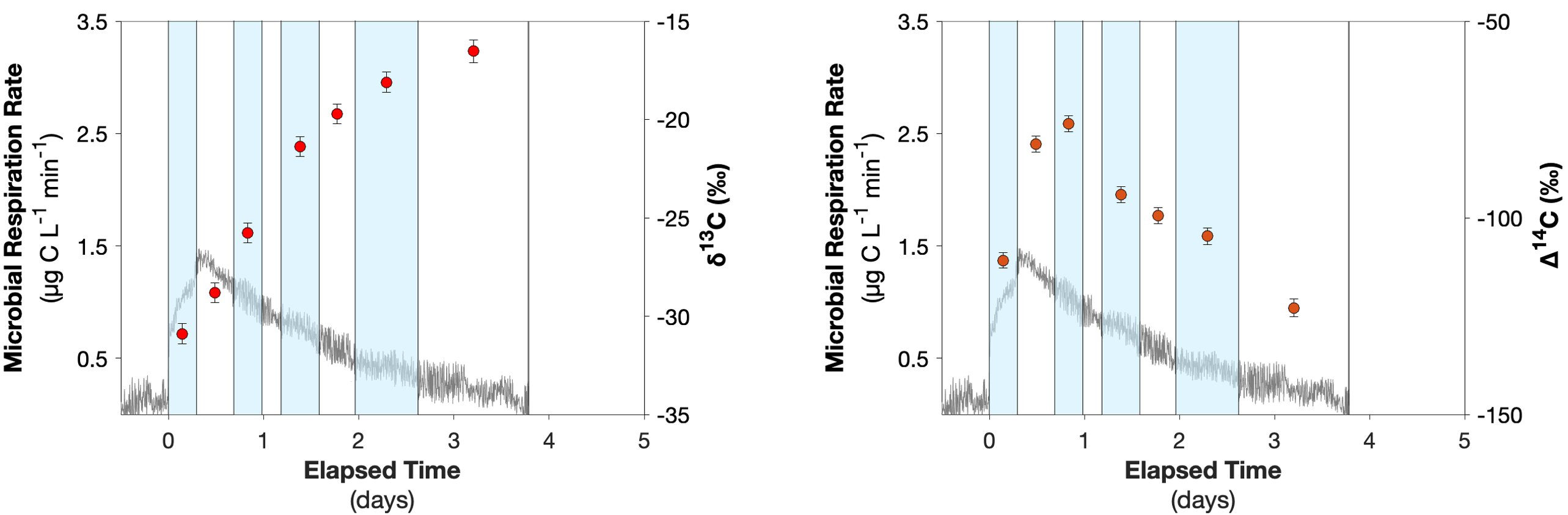

**Figure S3.** Microbial CO_2_ production rates (grey line), δ^13^C and Δ^14^C (circles) signatures of respired CO_2_ observed during co-culture incubation of *Vibrio splendidus* 1A01 and *Pseudoalteromonas* sp. 3D05 with (A) hydrothermal and (B) unimpacted Guaymas Basin sediment. The width of each box spans the time interval during which each CO_2_ fraction was collected for isotopic analysis, with the corresponding data point plotted at the mid-point for each fraction.

**A**

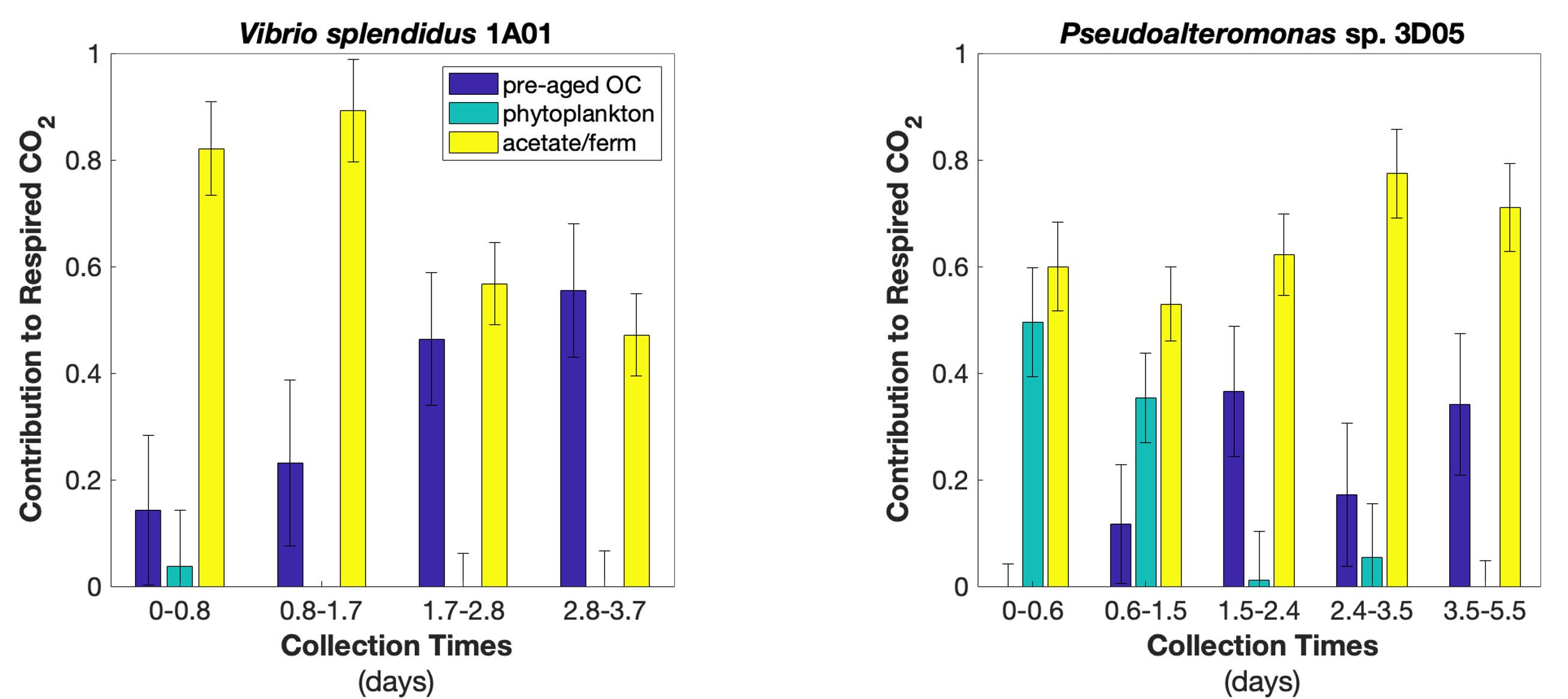

**B**

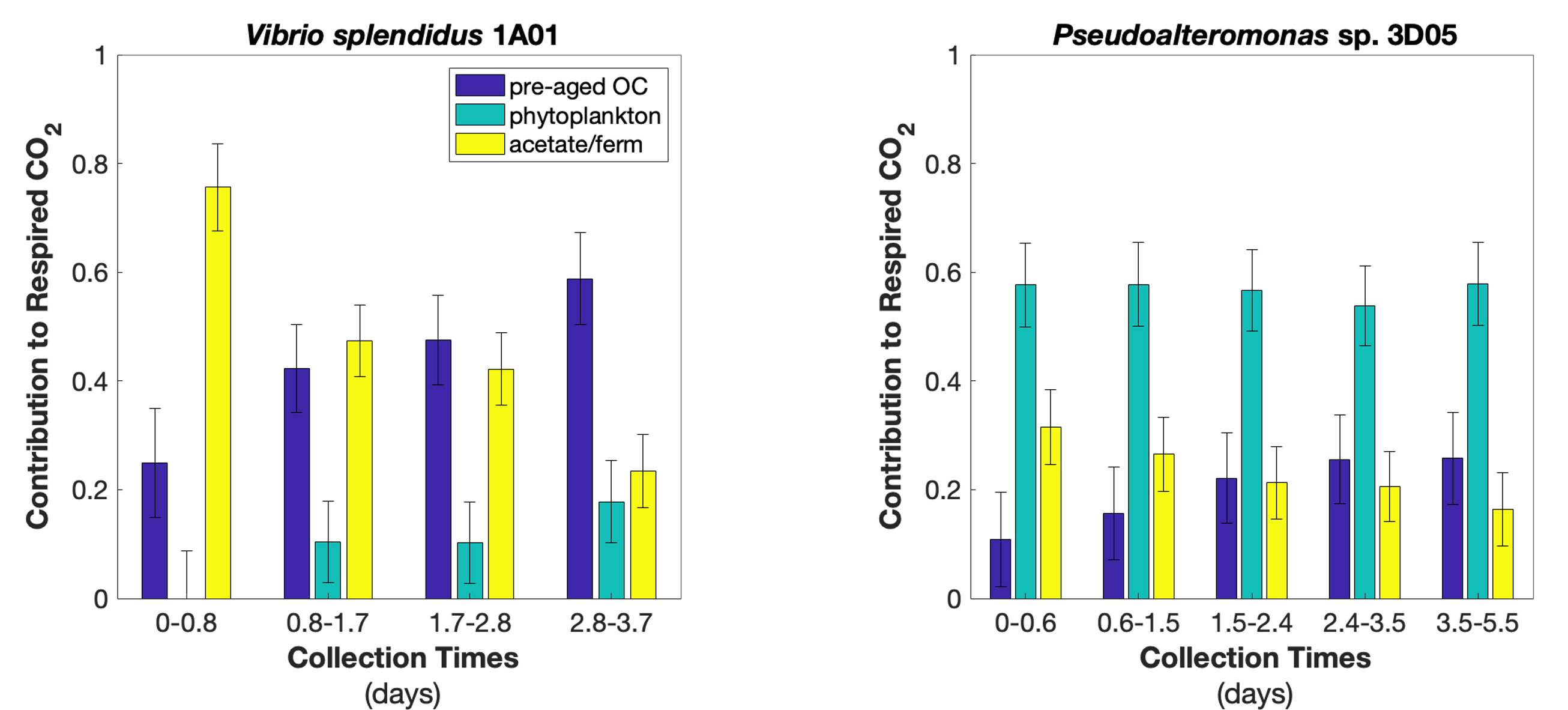

**Figure S4.** Differential utilization of carbon substrates between species. Estimated contributions of potential carbon sources to respired CO_2_ during individual incubations of *Vibrio splendidus* 1A01 and *Pseudoalteromonas* sp. 3D05 with (A) hydrothermal and (B) unimpacted Guaymas Basin sediment as described in Mahmoudi et al. (2020). Relative contributions were estimated using a three-end mass balance model. Percentages and uncertainties were estimated as means and standard deviations of solutions to 3 simultaneous mass-balance equations that were solved 10,000 times, in which normally distributed pseudo-random nose was added to each isotope ratio measurement and isotopic signature.

**A**

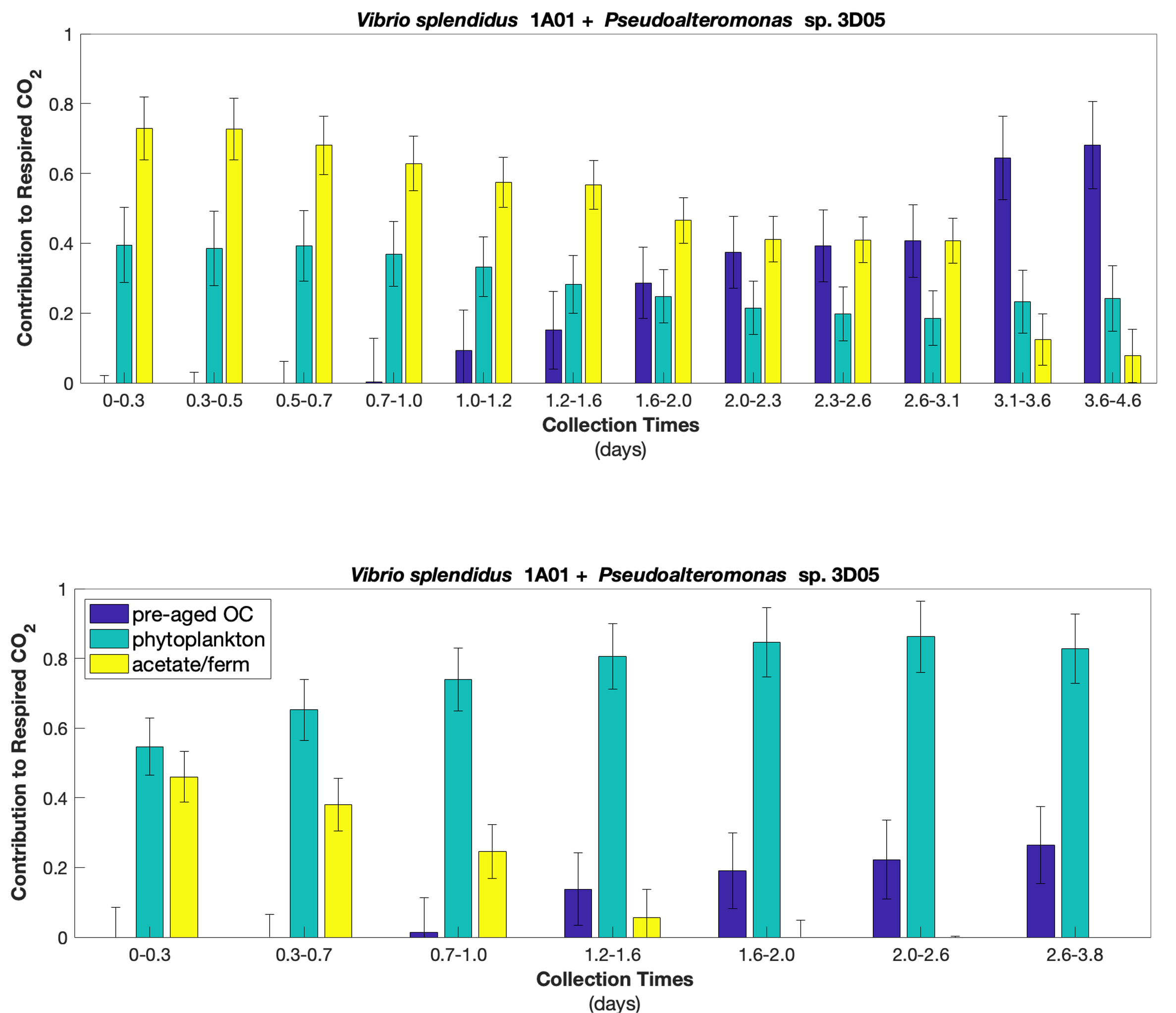

**B**

**Figure S5.** Differential utilization of carbon substrates between species. Estimated contributions of potential carbon sources to respired CO_2_ during co-culture incubations of *Vibrio splendidus* 1A01 and *Pseudoalteromonas* sp. 3D05 with (A) hydrothermal and (B) unimpacted Guaymas Basin sediment. Relative contributions were estimated using a three-end mass balance model. Percentages and uncertainties were estimated as means and standard deviations of solutions to 3 simultaneous mass-balance equations that were solved 10,000 times, in which normally distributed pseudo-random nose was added to each isotope ratio measurement and isotopic signature.

**Table S2.** Genome features of the isolates used for the competition experiments in this study. The genomes are deposited at NCBI under Bioproject # PRJNA414740 and the respective accession numbers below. The total copy number of genes encoding for known extracellular enzymes was determined from the annotated genomes and is described in Mahmoudi et al., (2020).

|  | ***Vibrio splendidus 1A01*** | ***Pseudoalteromonas sp. 3D05*** |
| --- | --- | --- |
| Genome size^1^ (MB) | 5.76 | 4.47 |
| GC content (%) | 44 | 40 |
| Coding density (genes per KB) | 0.90 | 0.90 |
| Number of coding genes | 5195 | 4040 |
| Accession nr. | PDUR00000000 | PDUS00000000 |
| *Gene copies for known extracellular enzymes* |  |  |
| Peptidase | 7 | 30 |
| Glucosidase | 0 | 1 |
| Chitinase | 5 | 7 |

^1^estimated from combined contig length

**Table S3.** Components added to make 2 L of modified Tibbles-Rawling minimal media [14].

| **Component** | **Amount** |
| --- | --- |
| **Part 1** | |
| NaCl | 51.9 g |
| MgSO4•7H2O | 6 g |
| MgCl2•6H2O | 4 g |
| CaCl2•2H2O | 0.24 g |
| Tris (1M; pH 8.0) | 50 mL |
| Na_2­_EDTA (0.5M) | 5.4 mL |
| NH4Cl (1M) | 20 mL |
| MilliQ water | 850 mL |
| **Part 2** | |
| K2HPO4 | 1.6 g |
| KH2PO4 | 0.4 g |
| MilliQ water | 850 mL |
| **Additives** |  |
| FeSO4 solution | 2 mL |
| Na2MoO4 solution | 2 mL |
| Trace metals (1000X) | 2 mL |
| Vitamins (1000X) | 2 mL |
| MilliQ water (total) | Fill to 2,000 mL |
| **1000X Vitamin Solution^1^** | **mg/L in MQ Water** |
| 4-aminobenzoate (4-aminobenzoic acid) | 40 mg |
| D(+)-Biotin | 10 mg |
| Folate (folic acid) | 30 mg |
| Lipoate ((±)-α-Lipoic acid) | 10 mg |
| Nicotinate (Nicotinic acid) | 100 mg |
| Ca-D-+-pantothenate | 50 mg |
| Pyridoxamine dihydrochloride | 100 mg |
| Thiamine hydrochloride | 100 mg |
| Vitamin B_12_ | 50 mg |
| **1000X Trace Metal Solution^2^** | **g/L in MQ Water** |
| H_3_BO_3_ | 2.86 g |
| MnCl_2_ • 4H_2_O | 1.81 g |
| ZnSO_4_ • 7H_2_O | 0.079 g |
| CuSO_4_ • 5H_2_O | 0.079 g |
| Co(NO_3_)_2_ • 6H_2_O | 0.0494 g |
| NiCl_2_ • 6H_2_O | 0.005 g |

^1^Finster et al., 1992 [15]

^2^Tibbles and Rawlings, 1994 [14]

**Table S4.** Details of hydrothermal and unimpacted Guaymas Basin sediment used in this study (adapted from Mahmoudi et al., 2020). Location of sampling stations (latitude and longitude in degrees, minutes, and decimal seconds), in-situ temperature range of sediment samples based on local temperature gradients in the sediment, total organic carbon (TOC) content and bulk isotopic composition of sediments.

|  | **Latitude** | **Longitude** | **Water depth (m)** | **In situ temp**  **(ºC)** | **Total Organic Carbon (TOC %)** | **δ^13^C (‰)** | **Δ^14^C (‰)** |
| --- | --- | --- | --- | --- | --- | --- | --- |
| Hydrothermal sediment | 27ºN02.70 | 111ºW23.10 | 2000 | 3-90ºC | 4.2 | -24.5 | -234 |
| Unimpacted sediment | 27ºN30.35 | 111ºW40.85 | 1725 | 3-5ºC | 4.0 | -24.9 | -226 |

**Table S5.** Details of primer sets selected for dPCR quantification of *Vibrio splendidus* 1A01 and *Pseudoalteromonas* sp. 3D05.

| **Target species** | **Target genomic region** | **Oligonucleotide binding site** | **Sequence (5’ – 3’)** | **Product Length (pb)** |
| --- | --- | --- | --- | --- |
| *Vibrio splendidus* 1A01 | CSB62_11365 segment | F927,797  R927,971  P 927,921 | GGATTATCAACCCACGCCC  CACGGTTTCATCTAGGGCC  6-FAM/CAA GTA CCA /ZEN/TCA CCG AGC ATT GAA CAG ACC /IABkFQ | 174 |
| *Pseudoaltermonas* sp. 3D05 | CSC79_13810 segment | F 109,787  R109,968  P 109,839 | CCTTACCGACTAATCCCGC  ATTACCCACTTGCCCGC  HEX/ACA ACC CAC /ZEN/CAC TCG CAT GAT GAT ACT GTC /IABkFQ | 181 |

F= Forward primer; R= Reverse primer; P= Probe

|  | **Fraction #** | **Collection Times (days)** | **Collection Duration (hr)** | **Mass CO_2_ (µg C)** | **δ^13^C (‰)** | **∆^14^C (‰)** |
| --- | --- | --- | --- | --- | --- | --- |
| **Hydrothermal sediment** |  |  |  |  |  |  |
| *Vibrio* sp. 1A01^1^ | 1 | 0.0 – 0.8 | 20.3 | 762 ± 3 | -40 ± 0.5 | -357 ± 2 |
|  | 2 | 0.8 – 1.7 | 20.3 | 532 ± 2 | -42 ± 0.5 | -418 ± 2 |
|  | 3 | 1.7 – 2.8 | 25.7 | 632 ± 3 | -35 ± 0.5 | -401 ± 2 |
|  | 4 | 2.8 – 3.7 | 23.0 | 416 ± 2 | -33 ± 0.6 | -405 ± 2 |
| *Pseudoalteromonas* sp. 3D05^1^ | 1 | 0.0 – 0.6 | 15.1 | 593 ± 3 | -34 ± 0.5 | -185 ± 2 |
|  | 2 | 0.6 – 1.5 | 19.7 | 1111 ± 5 | -33 ± 0.5 | -247 ± 2 |
|  | 3 | 1.5 – 2.4 | 23.9 | 1135 ± 5 | -36 ± 0.5 | -380 ± 2 |
|  | 4 | 2.4 – 3.5 | 25.7 | 989 ± 4 | -39 ± 0.5 | -353 ± 2 |
|  | 5 | 3.5 – 5.5 | 48.0 | 758 ± 3 | -38 ± 0.6 | -400 ± 2 |
| *Vibrio* sp. 1A01 + *Pseudoalteromonas* sp. 3D05 | 1 | 0.0-0.3 | 7.1 | 1310 ± 3 | -37 ± 0.5 | -218 ± 2 |
|  | 2 | 0.3-0.5 | 5.6 | 1845 ± 4 | -37 ± 0.5 | -222 ± 2 |
|  | 3 | 0.5-0.7 | 5.1 | 1679 ± 4 | -36 ± 0.5 | -222 ± 2 |
|  | 4 | 0.7-1.0 | 5.7 | 1799 ± 4 | -35 ± 0.5 | -235 ± 2 |
|  | 5 | 1.0-1.2 | 5.1 | 1354 ± 3 | -34 ± 0.5 | -253 ± 2 |
|  | 6 | 1.2-1.6 | 9.1 | 1676 ± 4 | -34 ± 0.5 | -274 ± 2 |
|  | 7 | 1.6-2.0 | 9.5 | 1631 ± 4 | -32 ± 0.5 | -294 ± 2 |
|  | 8 | 2.0-2.3 | 7.4 | 1003 ± 2 | -31 ± 0.5 | -311 ± 2 |
|  | 9 | 2.3-2.6 | 7.7 | 897 ± 2 | -31 ± 0.5 | -318 ± 2 |
|  | 10 | 2.6-3.1 | 11.3 | 1095 ± 2 | -31 ± 0.5 | -232 ± 2 |
|  | 11 | 3.1-3.6 | 13.3 | 418 ± 1 | -25 ± 0.5 | -322 ± 2 |
|  | 12 | 3.6-4.6 | 24.7 | 431 ± 1 | -24 ± 0.6 | -321 ± 2 |
| **Unimpacted sediment** |  |  |  |  |  |  |
| *Vibrio* sp. 1A01^1^ | 1 | 0.0 – 0.8 | 20.4 | 472 ± 2 | -39 ± 0.5 | -292 ± 2 |
|  | 2 | 0.8 – 1.7 | 19.9 | 570 ± 2 | -33 ± 0.5 | -314 ± 2 |
|  | 3 | 1.7 – 2.8 | 25.9 | 488 ± 2 | -32 ± 0.5 | -328 ± 2 |
|  | 4 | 2.8 – 3.7 | 23.7 | 380 ± 2 | -28 ± 0.6 | -341 ± 2 |
| *Pseudoalteromonas* sp. 3D05^1^ | 1 | 0.0 – 0.7 | 16.5 | 871 ± 4 | -28 ± 0.5 | -134 ± 2 |
|  | 2 | 0.7 – 1.6 | 21.5 | 629 ± 3 | -27 ± 0.5 | -146 ± 2 |
|  | 3 | 1.6 – 2.6 | 24.1 | 748 ± 3 | -26 ± 0.5 | -165 ± 2 |
|  | 4 | 2.6 – 3.6 | 25.3 | 519 ± 2 | -26 ± 0.5 | -180 ± 2 |
|  | 5 | 3.6 – 5.5 | 44.5 | 558 ± 2 | -25 ± 0.6 | -172 ± 2 |
| *Vibrio* sp. 1A01 + *Pseudoalteromonas* sp. 3D05 | 1 | 0.0-0.3 | 7.1 | 818 ± 2 | -30 ± 0.5 | -111 ± 2 |
|  | 2 | 0.3-0.7 | 9.4 | 1477 ± 3 | -29 ± 0.5 | -81 ± 2 |
|  | 3 | 0.7-1.0 | 7.1 | 908 ± 2 | -26 ± 0.5 | -76 ± 2 |
|  | 4 | 1.0-1.2 | 4.8 | 524 ± 1 | -- | -- |
|  | 5 | 1.2-1.6 | 9.6 | 710 ± 2 | -21 ± 0.5 | -94 ± 2 |
|  | 6 | 1.6-2.0 | 9.1 | 536 ± 1 | -20 ± 0.5 | -99 ± 2 |
|  | 7 | 2.0-2.6 | 15.8 | 615 ± 1 | -18 ± 0.5 | -105 ± 2 |
|  | 8 | 2.6-3.8 | 27.9 | 352 ± 1 | -17 ± 0.6 | -123 ± 2 |

**Table S6.** Masses and isotopic values of microbially-respired CO_2_ collected during co-culture incubation of *Vibrio splendidus* 1A01 and *Pseudoalteromonas* sp. 3D05 with Guaymas Basin sediment.

^1^Measured by Mahmoudi et al. (2020) during mono-culture incubations.
